# Captive-reared migratory monarch butterflies show natural orientation when released in the wild

**DOI:** 10.1101/2020.01.24.919027

**Authors:** Alana A. E. Wilcox, Amy E. M. Newman, Nigel E. Raine, D. Ryan Norris

## Abstract

Eastern North American migratory monarch butterflies (*Danaus plexippus*) have faced sharp declines over the last two decades. Although captive rearing has been used as an important tool for engaging the public and supplementing conservation efforts, a recent study that tested monarchs in a flight simulator suggested that captive-reared monarchs lose their capacity to orient southward during fall migration to their Mexican overwintering sites. We raised offspring of wild-caught monarchs on swamp milkweed (*Asclepias incarnata*) and, after eclosion, individuals were either tested in a flight simulator or radio-tracked in the wild using array of over 100 automated telemetry towers. While only 33% (7/39) of monarchs tested in the flight simulator showed strong southeast to southwest orientation, 97% (28/29) of the radio-tracked individuals were detected by automated towers south or southeast of the release site, up to 200 km away. Our results suggest that, though captive rearing of monarch butterflies may cause temporary disorientation, proper orientation is likely re-established after exposure to natural skylight cues.

## Introduction

Captive rearing and reintroduction of animals into the wild can be an effective tool for mitigating the decline of wild populations [1]. Capacity for acclimation in captivity varies among species [2-4], with some species, such as cheetahs (*Acinonyx jubatus*; [4-5]), being notoriously difficult to maintain or having lower fitness in captivity. Animal behaviour is known in differ between captive and wild populations of mammals [6], fish [7], and insects [8,9], and abnormal behaviour in captive populations of mammals is well documented [10-11]. However, only a single study, conducted on monarch butterflies (*Danaus plexippus*), has shown the potential for long-term behavioural impacts of captive-reared insects intended to be released in the wild [12].

In late fall, monarch butterflies undergo up to a 4,000 km migration from the mid-western and north-eastern United States and south-eastern Canada to Cerro Pelón and Sierra Madre Oriental mountains in Mexico [13-15]. Monarch butterflies are often reared in captivity by hobbyists and conservationists aiming to contribute towards population recovery of this species-at-risk. However, a recent study provided evidence that wild caught and chamber-reared monarch butterflies (i.e., reared from eggs indoors in autumn-like conditions until adult emergence (eclosion)) did not show normal southern orientation [12]. These results were obtained when individual adult butterflies were tested immediately after eclosion in a confined flight simulator that measured directional orientation. The authors concluded that the popular activity of hobbyists and conservationists in rearing captive monarchs for release would be an ineffective conservation practice to help boost migratory populations. However, the possibility remains that monarch butterflies released in the wild are able to show proper orientation if they can recalibrate their internal compass with exposure to natural skylight cues, an external cue known to be critical to the functioning of the molecular clock that governs directional flight [16].

In this study, we reared monarch butterflies in captivity and tested them in a confined flight simulator. Then, using an array of over 100 automated telemetry towers [17-18], butterflies reared in the same captive conditions were released and subsequently radio-tracked in the wild to evaluate whether captive-reared monarchs can reorient in a natural southward direction.

## Methods

This study was part of a larger project testing the effect of exposure to the neonicotinoid insecticide clothianidin on orientation of fall migratory monarch butterflies. Monarch butterflies were reared in captivity on swamp milkweed (*Asclepias incarnata*) grown in commercial soil either treated with 4, 8, 15 or 25 ng/g of clothianidin or an untreated control (see Supplementary Material). Despite these rearing conditions, there was no measurable effect of neonicotinoid exposure on orientation (results forthcoming). After eclosion, we tested the migratory orientation of adult monarchs in either a flight simulator or radio-tracked butterflies released into the wild.

### Flight simulator testing

For a subset of monarch butterflies (n = 54), we assessed monarch orientation during the fall migratory period (17-23 September 2018) using flight simulators (Figure 1a). Flight simulators were set up on the roof of the University of Guelph Phytotron (Guelph, ON) and arranged so that no surrounding buildings could obstruct the view of individuals while in the flight cylinder [19]. Tests occurred during daylight (9:30-16:00 EST) when the sun was fully visible from the simulator to ensure consistency of polarized light cues [19-20]. Individual butterflies were tethered to an L-shaped rod (modified to approximately 2.5 cm; CAT # 718000, 0.05 in x 15.2 in Tungsten Rods, A-M Systems, Washington, USA) inserted at the front of the dorsal thorax, avoiding flight muscle, and secured with super glue (All Purpose Krazy Glue No Run Gel, Elmer’s Products, High Point, NC, USA). Each tether was attached to a digital encoder that allowed 360° rotation and recorded orientation at 3° intervals [19]. The encoder was adhered to a plexiglass rod supported within a large cylinder and attached to a nearby computer to record directional data [19]. A fan at the base of the flight simulator provided airflow to encourage flight. Each monarch was flown once for 12 minutes, with the initial 2 minutes for acclimation and to minimize the impacts of stress-induced unidirectional flight response [21]. Monarchs were removed (n = 15) from the study if they did not show a characteristic pattern of flight (i.e., strong flapping with intermittent gliding).

**Figure 1.**
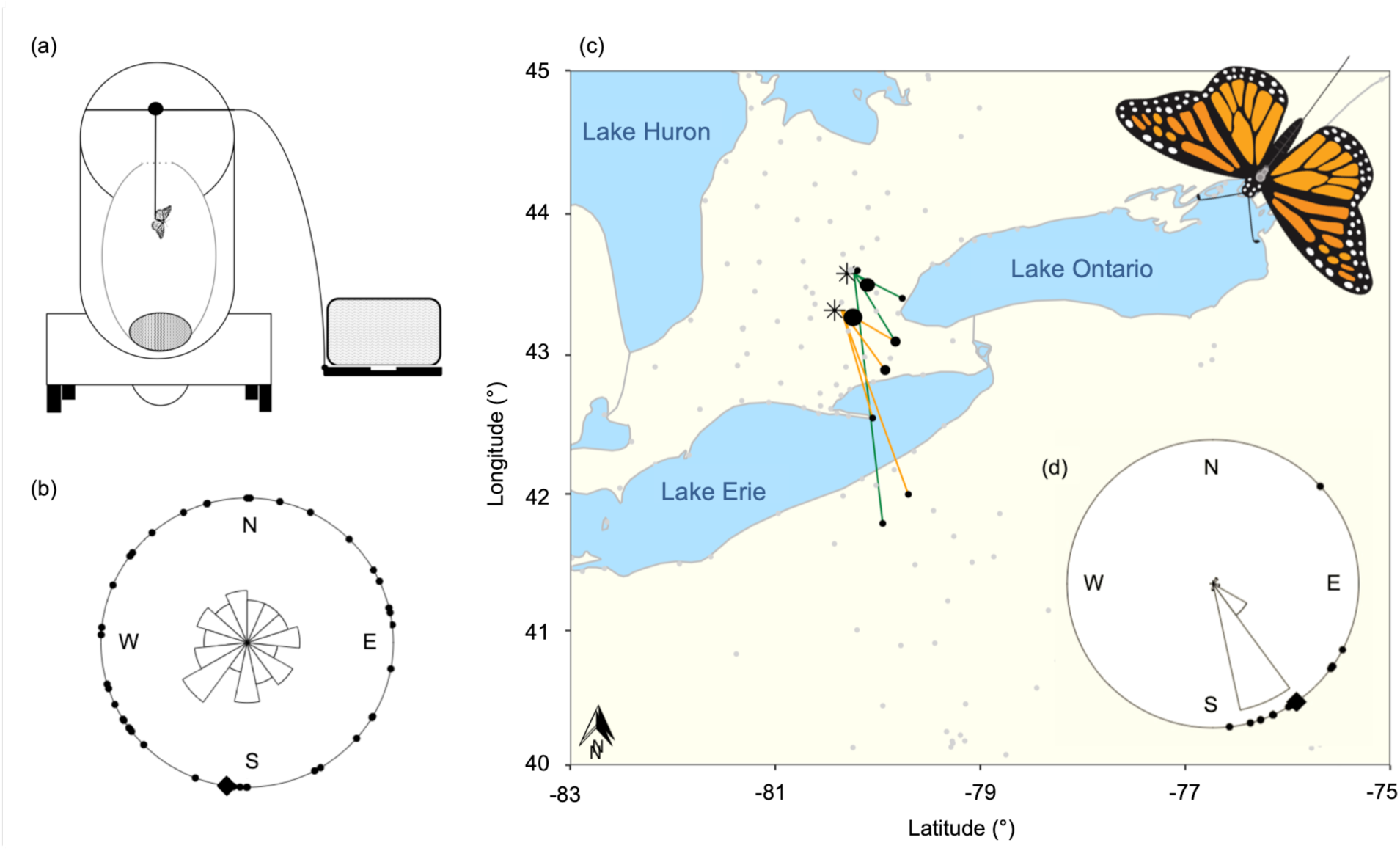
Orientation of captive-reared eastern North American migratory monarch butterflies (*Danaus plexippus*) (a) flown in a flight simulator for 10 minutes in Guelph, ON in September 2018. The (b) direction of flight (σ = 188°, n = 39; r = 0.30) of flight for individual monarchs (•) with the group mean direction (♦) is shown in a circular plot, where each section of the central windrose indicates the proportion of individuals with directional flight. Group mean direction is indicated as a solid line and each section of the windrose indicates the proportion of individuals with directional flight. (c) Map of the direction of flight for monarch butterflies released in Guelph, ON in September 2017 (n = 9, green lines) and Cambridge, ON in October 2018 (n = 20, orange lines). Symbols indicate the sites of release (✴) and location of first detection at a Motus tower (•), with the relative size referring to the number of detections at that tower (lowest number of detections at a tower = 1, highest number of detections at a tower = 5). Grey dots indicate Motus towers that were active at the time of releases. (d) Circular plot shows the direction of flight for radio-tracked monarch butterflies (σ = 145°, n = 29, r = 0.71).

### Radio-telemetry tracking

We tracked monarch butterflies using radio-transmitters during early migration. Monarchs were outfitted with 200 mg NanoTags (Lotek Wireless Fish & Wildlife Monitoring, Newmarket, ON, Canada), each programmed to emit unique 166.380 MHz pulses every 1.5 seconds to maximize the probability of detection and allow individual identification [18]. Large individuals (>0.3 g) were selected to minimize weight restrictions imposed by the tags and maximize the capacity for long distance flight. On 5 October 2017, 41 monarch butterflies were released in an open field in Guelph, ON (43.6°N, -80.2°W), centered between adjacent Motus towers. On 27 September 2018, 43 monarchs were released on a hill, above tree line, at the base of the Cambridge RARE Motus tower (43.4°N, −80.4°W) in Cambridge, ON. The Motus telemetry array consists of more than 100 independent VHF telemetry towers across southern Ontario and the northern United States, with towers in all directions around the site of release [17-18]. False detections were removed from analysis following the procedures outlined by Crewe et al. [22]. We ran preliminary filters to remove detections with run lengths (i.e., number of detections) <2 and false detections as a result of noise (e.g., detections prior to release or beyond the species range, towers recording spurious detections). We also examined ambiguous detections manually, using contextual information to identify true detections [22]; for instance, removing detections that bounced between multiple towers and/or countries. We also removed detections recorded on the day of release at adjacent towers with overlapping detection ranges to the site of release to avoid inaccurately assigning a direction of flight when the monarchs had not left the area. This resulted in true detections for 9 monarch butterflies in 2017 and 20 monarchs in 2018.

### Statistical analysis

North American migratory monarch butterflies orient in a southward direction when flown in a flight simulator [23-25]. We calculated the mean direction (0° to 359°) and vector strength (r: 0 – 1) for each monarch butterfly flight using R version 3.4.1 [26]. Vector strength is a measure of concentration for circular data with high values indicating a tighter grouping around the mean direction [19,27].

To compare orientation between monarchs flown in the flight simulator and tracked using the Motus Wildlife Tracking System, we followed previously published procedures from Tenger-Trolander et al. [12]. We calculated the group mean direction, weighted by the group vector strength and each individual’s vector strength. For radio-tracked monarchs, the same weight was given to the direction flown by individual monarchs since the direction for each individual was recorded only once from release to the first detection at a Motus tower. In a separate analysis, we used a Rayleigh test in the R circular package to determine if the sample of monarchs flown in the flight simulator and tracked using the Motus telemetry array showed directional flight. Finally, we calculated the percentage of individual monarch butterflies that flew in the southward direction (i.e., in the southeast to southwest direction) and calculated Spearman’s Rank correlation coefficients to assess the relationship between distance travelled with time (i.e., greater distance travelled with a longer duration of time since release).

## Results

Although the mean direction for monarchs flown in the flight simulator was σ = 188° (SSW), individuals showed strong orientation in a variety of directions, including N, NW, NE, resulting in the sample only being weakly concentrated around the mean (Rayleigh test, n = 39, r = 0.30, p = 0.03; Figure 1b). Overall, only 33% of monarchs tested in the flight simulator oriented in the southeast to southwest direction (Supplementary Material Table S1). In contrast, 97% of radio-tracked monarchs (28/29) flew south to southeast (σ = 145°; Figure 1c-d; Supplementary Material Table S1). The direction of flight for radio-tracked monarchs was strongly concentrated around the mean (Rayleigh test, n = 29, r = 0.71, p < 0.001; Figure 1d). Monarchs were first detected 1 to 16 days after release (Supplementary Material Table S2) at towers from 11 km (52%, 15/29) up to 200 km (3%, 1/29) from the site of release and the number of days to first detection was correlated with distance from the release site (Spearman’s rank correlation, n = 29, r_s_= 0.70, p < 0.001).

## Discussion

Our results provide evidence that monarch butterflies raised in captivity, but later exposed to natural conditions (i.e., sunlight and photoperiod), can reset the mechanism governing directional flight, allowing them to properly orient southward towards Mexico after they are released into the wild. Monarch butterflies tested in the flight simulator generally oriented in the southward direction, but the data were widely distributed in all directions and only 33% flew in the southeast to southwest direction. When released into natural conditions 97% of monarchs flew in the south to southeast direction. Thus, while our study confirms the results from Tenger-Trolander et al. [12] that most captive-reared monarchs tested in a flight simulator do not show proper orientation towards their Mexican overwintering grounds, we also show that monarchs released in the wild are capable of recalibrating the mechanism responsible for directional flight. Therefore, we provide strong support for the practice of captive rearing as a conservation tool to supplement populations of monarch butterflies and improve recovery of this species-at-risk.

The results of our experiment suggest that outdoor environmental conditions are required for proper directional flight during migration. In monarchs, sunlight is received and processed by a molecular clock in the antennae [28-30]. Disruption of this molecular mechanism by restricting natural light results in disoriented flight [28], providing evidence that sunlight is required for monarchs to calibrate flight orientation. A similar recalibration with environmental cues was found in *Catharus* thrushes [31]. After exposure to experimental magnetic fields, Gray-cheeked thrushes (*C. minimus*) and Swainson’s thrush (*C. ustulatus*) were released and their flight patterns tracked using radio transmitters [31]. On the first night, birds flew westward, but corrected their orientation by the second night after they were exposed to ‘normal’ twilight cues and flew in the proper northward direction [31]. Though mechanisms underpinning flight orientation differ between birds and insects [32], it is likely that monarchs can also recalibrate the direction of flight using information obtained via skylight and other natural cues.

Captive rearing of monarch butterflies for wildlife education, commercial breeding programs or by hobbyists can enhance conservation efforts if precautions are taken to rear monarchs in conditions that allow exposure to natural environmental conditions. Though commercially reared monarchs tested by Tenger-Trolander et al. [12] showed a random orientation, the authors contrast their findings with a successful tag and release by Maeckle [33] where released monarchs were re-sighted in Mexico and our results clearly demonstrate that upon release monarchs regain proper orientation. We suggest that under proper rearing conditions, particularly exposure to sunlight, loss of orientation capacity may be negligible and future studies should determine the minimum duration of sunlight required to establish southward directional flight. Though the practice of captive rearing is contentious due to the potential for disease transmission [12,34-35] and concerns around genetic viability [12,34-36], when these risks are minimized, reintroduction of monarch butterflies to the wild could contribute towards reversing declines of migratory populations. Captive rearing of monarchs is not only a tool for conservation, but is also an extraordinary educational opportunity for the public to interact with nature and engage in conservation. The incredible social appeal of monarch butterflies and captive rearing for educational purposes encourages interactions between the public, educators, and scientists [37]. Thus, under proper conditions, captive rearing offers an opportunity for the public to engage in the conservation of this beloved and iconic species.

Although our results suggest that sunlight reestablishes southward directional flight in a North American fall migratory population of monarch butterflies, our experimental design did not allow us to investigate the duration of exposure to solar cues required for recalibration of the molecular clock mechanism. Nor were we able to test individual monarch butterflies in the flight simulator and then release the same individuals in the wild. Monarchs tested in the flight simulator were temporarily compromised due to the insertion of a rod into the front of the dorsal thorax and showed visible signs of exhaustion (i.e., lethargy) after testing. With the continued development of tracking technology, it is likely that we will soon have the ability to track monarchs and other insects at finer spatial resolutions and over multiple days during their migratory journey. When that occurs, our understanding of the proximate mechanisms that govern orientation and effects of captive rearing will likely improve.

Eastern North American monarch butterflies have undergone declines of over 80% in the last two decades [38]. These astonishing declines serve as a reminder of the challenges faced in conserving biodiversity, particularly of insects, and in the conservation of this species-at-risk. Moreover, with increasing awareness of numerous threats to monarch butterfly populations [38], extensive support has been garnered across Canada, the US, and Mexico for monarch conservation. Our results confirm studies on the impact of captive rearing on monarch butterflies [12], but also strongly contrasts previously published research [12]. Captive-reared monarchs regain proper flight orientation when released into the wild, demonstrating that the popular activity of rearing monarch butterflies from caterpillars in captivity can be a viable conservation tool and important education element to conserve this iconic pollinator species.

## Supporting information

Wilcox et al 2020 Supplementary Material

## Ethics

No ethical approval was required prior to conducting research. Research was conducted under an Ontario Ministry for Natural Resources Wildlife Scientific Collectors Permit (2017: #1086793; 2018: #1090000).

## Data accessibility

Motus data is available at https://motus.org/data/downloads (Project ID # 209).

## Author contributions

A.A.E.W., A.E.M.N., N.E.R. and D.R.N. conceived and designed the project. A.A.E.W. conducted the experimental work, analysed the data, and drafted the original manuscript. All authors contributed to writing and revising the manuscript.

## Acknowledgements

We would like to thank Taylor Van Belleghem, Angela Demarse, and Samantha Knight, as well as our team of volunteers, for assistance with data collection. Thank you to Mike Mucci and Tannis Slimmon for technical support and coordinating use of the University of Guelph Phytotron.

## References

1. D. G. Hughes, P. M. Bennett, Captive breeding and the conservation of invertebrates. Int. Zoo Yearb. 30, 45–51 (1991).

2. A. S. Chamove, G. S. Hosey, P. Schaetzel, Visitors excite primates in zoos. Zoo Biol. 7, 359–369 (1988).

3. C. Mettke, “Ecology and environmental enrichment – the example of parrots” in Research and Captive Propagation, U. Gansloßer, J. K. Hodges, W. Kaumanns, eds. (Filander Verlag, 1995).

4. G. J. Mason, Species differences in responses to captivity: stress, welfare and the comparative method. Trends Ecol. Evol. 25, 713–721 (2010).

5. K. Bauman, E. Blumer, A. Crosier, J. Fallon, G. Geise, J. Grisham, J. Ivy, S. Long, A. Rogers, K. Schwartz, K. Snodgass, E. Spevak, Global cheetah *ex situ* planning: linking managed populations working group. Conserv. Breed. Specialist Group News 21, 1–4 (2010).

6. R. J. Blanchard, K. J. Flannelly, D. C. Blanchard. Defensive behaviors of laboratory and wild *Rattus norvegicus*. J. Comp. Psychol. 100, 101–107 (1986).

7. A. Gro Vea Salvanes, V. Braithwaite. The need to understand the behaviour of fish reared for mariculture or restocking. ICES Journal of Marine Science 63, 345–354 (2006).

8. D. N. Fisher, A. James, R. Rodríguez-Muñoz, T. Tregenza. Behaviour in captivity predicts some aspects of natural behaviour, but not others, in a wild cricket population. Proc. Roy. Soc. B – Biol. Sci. 282, 20150708 (2015).

9. T. C. Ings, N. E. Raine, L. Chittka, A population comparison of the strength and persistence of innate colour preference and learning speed in the bumblebee *Bombus terrestris*. Behav. Ecol. Sociobiol. 63, 1207-1218 (2009).

10. L. P. Birkett, N. E. Newton-Fisher, How abnormal is the behaviour of captive, zoo-living chimpanzees? PLoS ONE 6, e20101 (2011).

11. M. E. McPhee, Generations in captivity increases behavioral variance: considerations for captive breeding and reintroduction programs. Biol. Conserv. 115, 71–77. (2004).

12. A. Tenger-Trolander, W. Lu, M. Noyes, M. R. Kronfrost, Contemporary loss of migration in monarch butterflies. Proc. Nat. Acad. Sci. USA 116, 14671–14676 (2019).

13. L. P. Brower, Understanding and misunderstanding the migration of the monarch butterfly (Nymphalidae) in North America: 1857–1995. J. Lepid. Soc. 49, 304–385 (1995)

14. F. A. Urquhart, “Migration” in The Monarch Butterfly (University of Toronto Press, 1960), pp. 77–94.

15. F. A. Urquhart, N. R. Urquhart, Autumnal migration routes of the eastern population of the monarch butterfly (*Danaus p. plexippus* L.; Danaidea; Lepidoptera) in North America to the overwintering site in the Neovolcanic Plateau of Mexico. Can. J. Zool. 56, 1759–1764 (1978).

16. S. M. Reppert, R. J. Gegear, C. Merlin, Navigational mechanism of migrating monarch butterflies. Trends Neurosci. 33, 399–406 (2010).

17. Motus (Motus Wildlife Tracking System), About Motus. Available at http://motus.org/about (2017).

18. P. D. Taylor, T. L. Crewe, S. A. Mackenzie, D. Lepage, Y. Aubry, Z. Crysler, G. Finney, C. M. Francis, C. G. Guglielmo, D. J. Hamilton, R. L. Holberton, P. H. Loring, G. W. Mitchell, D. R. Norris, J. Paquet, R. A. Ronconi, J. Smetzer, P. A. Smith, L. J. Welch, B. K. Woodworth, The Motus Wildlife Tracking System: a collaborative research network to enhance the understanding of wildlife movement. Avian Conserv. Ecol. 12, 8 (2017).

19. H. Mouritsen, R. Derbyshire, J. Stalleicken, O. Ø. Mouritsen, B. J. Frost, D. R. Norris, An experimental displacement and over 50 years of tag-recoveries show that monarch butterflies are not true navigators. Proc. Nat. Acad. Sci. USA 110, 7348–7353 (2013).

20. S. M. Reppert, H. Zhu, R. H. White, Polarized light helps monarch butterflies navigate. Curr. Biol. 14, 155–158 (2004).

21. S. M. Perez, O. R. Taylor, R. Jander, The effect of a strong magnetic field on monarch butterfly (*Danaus plexippus*) migratory behaviour. Naturwissenschaften 86, 140–143 (1999).

22. T. L. Crewe, Z. Crysler, P. Taylor, “Data cleaning” in Motus R Book: A Walk Through the Use of R for Motus Automated Radio-Telemetry Data. Available at https://motus.org/MotusRBook/.

23. O. Froy, A. L. Gotter, A. L. Casselman, S. M. Reppert, Illuminating the circadian clock in monarch butterfly migration. Science 300, 1303–1305 (2003).

24. P. A. Guerra, S. M. Reppert, Coldness triggers northward flight in remigrant monarch butterflies. Curr. Biol. 23, 419–423 (2013).

25. H. Mouritsen, B. J. Frost. Virtual migration in tethered flying monarch butterflies reveals their orientation mechanisms. Proc. Nat. Acad. Sci. USA 99, 10162–10166 (2012).

26. R Core Team, R: a language and environment for statistical computing. R Foundation for Statistical Computing, Vienna, Austria. Available at https://www.r-project.org/ (2015).

27. A. Pewsey, M. Neuhäuser, G. D. Ruxton, “Circular summary statistics” in Circular Statistics in R (Oxford University Press, 2013), pp. 21–34.

28. P. A. Guerra, C. Merlin, R. J. Gegear, S. M. Reppert, Discordant timing between antennae disrupts sun compass orientation in migratory monarch butterflies. Nat. Comm. 17, 958 (2012).

29. S. M. Reppert, D. R. Weaver, Coordination of circadian timing in mammals. Nature 418, 935–941 (2002).

30. S. M. Reppert. A colorful model of the circadian clock. Cell 124, 233–236 (2006).

31. W. W. Cochran, H. Mouritsen, M. Wikelski. Migrating songbirds recalibrate their magnetic compass daily from twilight cues. Science 304, 405–408 (2004).

32. H. Mouritsen, Long-distance navigation and magnetoreception in migratory animals. Nature 558, 50–59 (2018).

33. M. Maeckle, Five monarch butterflies tagged and released at San Antonio Festival made it to Mexico. Texas Butterfly Ranch. Available at https://texasbutterflyranch.com/2018/04/25/five-monarch-butterflies-tagged-and-released-at-san-antonio-festival-made-it-to-mexico/ (2018).

34. Journey North, Captive breeding and releasing monarchs. Available at https://journeynorth.org/tm/monarch/conservation_action_release.pdf (2015).

35. Monarch Joint Venture, Raising monarchs: why or why not? (Revised Handout). Available at https://monarchjointventure.org/news-events/news/revised-handout-raising-monarchs-why-or-why-not (2018).

36. J. R. Willoughby, J. A. Ivy, R. C. Lacy, J. M. Doyle, J. A. DeWoody. Inbreeding and selection shape genomic diversity in captive populations: implications for the conservation of endangered species. PLoS ONE 12, e0175996 (2017).

37. K. M. Gustafsson, A. A. Agrawal, B. V. Lewinstein, S. A. Wolf. The monarch butterfly through time and space: the social construction of an icon. BioSci. 65, 612–622 (2015).

38. W. E. Thogmartin, R. Wiederholt, K. Oberhauser, R. G. Drum, J. E. Diffendorfer, S. Altizer, O. R. Taylor, J. Pleasants, D. Semmens, B. Semmens, R. Erickson, K. Libby, L. Lopez-Hoffman. Monarch butterfly population decline in North America: identifying the threatening processes. Roy. Soc. Open Sci. 4, 170760 (2017).

