## Supplementary material for "Captive-reared migratory monarch butterflies show natural orientation when released in the wild": Wilcox et al 2020 Supplementary Material

Alana A. E. Wilcox[1*], Amy E. M. Newman[1], Nigel, E. Raine[2], and D. Ryan Norris[1,3]

[1] *Department of Integrative Biology, University of Guelph, Guelph, ON, N1G 2W1, Canada*

[2] *School of Environmental Sciences, University of Guelph, Guelph, ON, N1G 2W1, Canada*

[3] *Nature Conservancy of Canada, 245 Eglington Avenue East, Toronto, ON, M4P 3J1, Canada*

### Methods

#### *Milkweed*

Swamp milkweed (*Asclepias incarnata*) was grown in commercial soil (LA4 Sunshine Loosefill, Sungro Horticulture, Massachusetts, USA) treated with either at 4, 8, 15 or 25 ng/g of clothianidin (neonicotinoid insecticide) or a control (i.e., distilled water) with 4 plants per 6 square inch/1.68 L pot in environmental chambers at the University of Guelph Phytotron. Milkweed was watered twice daily until the soil was saturated. Room temperature was set at 29°C during the day and 23°C at night, 500 mol light (18L:6D) as outlined in Flockhart et al. [1]. Humidity was monitored hourly using a handheld thermohygrometer (Vaisala MI70 Measurement Indicator with HMP75 Humidity and Temperature Probe, Vasiala, Helsinki, Finland) with an average 77% (SD ±10%) RH. Plants were watered daily with reverse osmosis water and fertilized weekly with Plant-Prod Solutions fertilizer 17:5:17 NPK (Master Plant-Prod Inc., Brampton, ON, Canada). *Amblyseius swirskii* were introduced as a biocontrol (Bioline AgroSciences Swirskiline Biocontrol Agent and Biobest Swirskii-Breeding-System) measure to reduce the impact of thrips (Thysanoptera) [1].

#### *Capture and maintenance*

We raised monarch caterpillars from eggs laid by wild monarchs obtained from Gowanstown, ON (43.52°N, -81.08°W; male, n = 7; female, n = 13) on 14 August 2017 and the Guelph Lake Conservation Area (43.61°N, -80.26°W; male, n = 7; female, n = 11) from 2-6 August 2018. Wild monarch butterflies were held in coin envelopes (6.35 cm x 10.8 cm) inside an animal carrier kept at ambient temperature and humidity maintained with a damp cloth at the bottom of the carrier to avoid the wings drying out during transport to the University of Guelph. Butterflies were weighed (Denver Instrument PI-602 scale, Denver Instrument, Bohemia, NY, USA) to the nearest 0.01 g and individually hand-fed a 10% honey-water solution daily until satiation. Wild monarchs were mated in large mesh enclosures (60 cm height x 60 cm depth x 60 cm width) inside an incubator set at temperatures fluctuating between 29°C and 23°C with 500 mol light (18L:6D) and an average 77% (SD ±10%) RH. Enclosures contained untreated milkweed (i.e., grown in soil dosed with reverse osmosis water) and an artificial nectar source (i.e., 10% sucrose water) provided *ad libitum*.

We collected 192 eggs each year (n = 64 per treatment) by gently pressing a fine-tipped paintbrush along the edge of the egg and transferring to a milkweed leaf with residual latex holding the egg in place. In 2017, monarch caterpillars were reared directly on the milkweed plants with pots enclosed with finely perforated mosquito netting (Bulk Mosquito Netting, CAT # 09A04.73, Lee Valley, Ottawa, ON, Canada). Light, temperature and humidity in the University of Guelph Phytotron were maintained according to ambient conditions during the early fall in Guelph (43.5°N, -80.2°W; 13 hours light: 11 hours dark) at 21°C day:11°C night and an average 87% (SD ±6%) RH. Caterpillars were fed milkweed *ad libitum* until pupation when chrysalids were then transferred to mesh enclosures (60 cm height x 60 cm depth x 60 cm width) in the University of Guelph Phytotron separated by treatment after eclosion from 19 September – 3 October 2017. In 2018, leaves with eggs were placed in large plastic containers and enclosed with finely perforated mosquito netting. Environmental conditions were replicated from those used in 2017, except that chrysalids were transferred to mesh enclosures (120 cm x 120 cm x 120 cm; Popadome Plant Dome, CAT # XC515, Lee Valley, Ottawa, ON, Canada) in the laboratory where lighting cycle was variable, but supplemented by negligible foyer lighting. Eclosion occurred from 14-19 September 2018. Adult monarchs were hand-fed daily and provided dishes with a sucrose solution within the enclosures [1]. We examined each individual for *Ophryocystis elektroscirrha* protozoan parasites by applying clear tape to the abdomen and analyzing tape for spores under a microscope at 400x [2] and infected butterflies were removed from the study. All procedures will be conducted under the Ontario Ministry for Natural Resources Wildlife Scientific Collectors Permit (2017: #1086793; 2018: #1090000).

**References**

1. D. T. T. Flockhart, T. G. Martin, D. R. Norris, Experimental examination of intraspecific density-dependent competition during the breeding period in monarch butterflies (*Danaus plexippus*). *PLoS ONE* **7**, e45080 (2012).

2. S. M. Altizer, K. S. Oberhauser. Effects of the protozoan parasite *Ophryocystis elektroscirrha* on the fitness of monarch butterflies (*Danaus plexippus). J. Invert. Pathol.* **74**, 76–88 (1999).

**Supplementary Tables**

**Table S1.** Eastern North American migratory monarch butterflies (*Danaus plexippus*) reared in environmental chambers simulating autumn conditions (day: 29°C, night 23°C and 77% RH) until pupation. Monarchs (n = 39) were tested in an outdoor flight simulator that recorded orientation between 0° to 359°. The mean direction (° ± SD) and vector strength (r), representing the concentration of the data between 0 (data are evenly spread) to 1 (data are concentrated around the mean), was calculated for each individual. Rayleigh test was used to determine whether monarchs showed directional flight ($p < 0.05$). Each row represents an individual monarch butterfly.

| **Flight simulator** | | | | | |
|---|---|---|---|---|---|
| | | | | **Rayleigh test** | |
| **Cardinal direction** | **Mean direction (°)** | **SD (°)** | **r** | **z** | **p** |
| N | 0 | 0 | 1.00 | 1.00 | < 0.001 |
| N | 1.16 | 2.66 | 0.05 | 0.05 | 0.001 |
| N | 12.95 | 2.21 | 0.78 | 0.78 | < 0.001 |
| NNE | 25.70 | 2.50 | 0.29 | 0.29 | < 0.001 |
| NE | 44.37 | 2.86 | 0.04 | 0.04 | < 0.001 |
| ENE | 59.88 | 2.53 | 0.10 | 0.09 | < 0.001 |
| ENE | 64.89 | 2.48 | 0.03 | 0.03 | 0.09 |
| ENE | 76.11 | 2.03 | 0.27 | 0.27 | < 0.001 |
| ENE | 78.00 | 2.58 | 0.38 | 0.38 | < 0.001 |
| E | 82.94 | 2.65 | 0.76 | 0.76 | < 0.001 |
| E | 100.43 | 1.21 | 1.00 | 0.99 | < 0.001 |

| | | | | | |
|---|---|---|---|---|---|
| ESE | 120.66 | 2.16 | 0.61 | 0.61 | < 0.001 |
| SE | 121.15 | 2.23 | 0.32 | 0.32 | < 0.001 |
| SE | 131.45 | 1.21 | 1.00 | 0.99 | < 0.001 |
| SSE | 149.85 | 1.78 | 0.66 | 0.66 | < 0.001 |
| SSE | 152.47 | 2.26 | 0.68 | 0.68 | < 0.001 |
| S | 179.98 | 2.35 | 0.85 | 0.85 | < 0.001 |
| S | 180.21 | 2.98 | 0.12 | 0.12 | < 0.001 |
| S | 182.74 | 2.44 | 0.82 | 0.82 | < 0.001 |
| S | 185.31 | 1.78 | 0.87 | 0.87 | < 0.001 |
| SSW | 200.73 | 2.07 | 0.5 | 0.50 | < 0.001 |
| SW | 225.00 | 1.50 | 0.93 | 0.93 | < 0.001 |
| SW | 232.13 | 2.40 | 0.29 | 0.29 | < 0.001 |
| SW | 233.28 | 1.39 | 0.42 | 0.42 | < 0.001 |
| SW | 233.89 | 1.60 | 0.62 | 0.62 | < 0.001 |
| WSW | 237.42 | 2.89 | 0.37 | 0.37 | < 0.001 |
| WSW | 238.00 | 2.14 | 0.43 | 0.43 | < 0.001 |
| WSW | 244.67 | 2.65 | 0.74 | 0.74 | < 0.001 |
| WSW | 251.47 | 2.51 | 0.19 | 0.19 | < 0.001 |
| WSW | 253.29 | 0.83 | 0.75 | 0.75 | < 0.001 |
| W | 273.14 | 2.70 | 0.61 | 0.61 | < 0.001 |
| W | 276.06 | 2.30 | 0.42 | 0.42 | < 0.001 |
| WNW | 293.41 | 2.69 | 0.41 | 0.41 | < 0.001 |
| WNW | 306.76 | 1.71 | 0.29 | 0.29 | < 0.001 |
| NW | 308.41 | 2.88 | 0.25 | 0.25 | < 0.001 |
| NW | 319.48 | 1.09 | 0.97 | 0.97 | < 0.001 |

| | | | | | |
|---|---|---|---|---|---|
| NNW | 334.23 | 0.80 | 0.98 | 0.98 | < 0.001 |
| NNW | 343.97 | 1.85 | 0.76 | 0.76 | < 0.001 |
| NNW | 344.26 | 1.33 | 0.78 | 0.78 | < 0.001 |

**Table S2.** Eastern North American migratory monarch butterflies (*Danaus plexippus*) reared in environmental chambers simulating autumn conditions (day: 29°C, night 23°C and 77% RH) until pupation. Monarch butterflies (n = 29) were then released with radio-telemetry tags and tracked using the Motus telemetry array. The number of days and distance to first detection, as well as the direction (°) of flight after release in Guelph (2017) and Cambridge, ON (2018) was recorded. Each row represents an individual monarch butterfly.

| | Telemetry tracking | | | |
|---|---|---|---|---|
| **Year** | **No. days after release** | **Distance (km)** | **Cardinal direction** | **Direction (°)** |
| 2017 | 4 | 47 | NE | 47.49 |
| | 3 | 43 | ESE | 117.24 |
| | 3 | 14 | SE | 124.92 |
| | 3 | 14 | SE | 124.92 |
| | 3 | 14 | SE | 124.92 |
| | 3 | 14 | SE | 124.92 |
| | 3 | 14 | SE | 124.92 |
| | 5 | 63 | SSE | 148.80 |
| | 16 | 20 | S | 173.44 |
| 2018 | 3 | 52 | SE | 126.10 |
| | 8 | 64 | SSE | 147.3 |
| | 7 | 64 | SSE | 155.63 |
| | 1 | 12 | SSE | 155.63 |
| | 2 | 52 | SSE | 155.63 |
| | 1 | 64 | SSE | 155.63 |
| | 2 | 52 | SSE | 155.63 |

| | | | |
|---|---|---|---|
| 2 | 52 | SSE | 155.63 |
| 1 | 12 | SSE | 155.63 |
| 1 | 12 | SSE | 155.63 |
| 1 | 12 | SSE | 155.63 |
| 2 | 12 | SSE | 155.63 |
| 1 | 12 | SSE | 155.63 |
| 7 | 12 | SSE | 155.63 |
| 2 | 12 | SSE | 155.63 |
| 2 | 12 | SSE | 155.63 |
| 1 | 12 | SSE | 155.63 |
| 8 | 162 | SSE | 160.88 |
| 8 | 64 | SSE | 160.88 |
| 8 | 95 | SSE | 165.07 |
